# Anti-diabetic Potential of Aqueous Extract of *Moringa oleifera, Ocimum gratissimum* and *Vernonia amygdalina* in Alloxan-Induced Diabetic Rats

**DOI:** 10.1101/2020.08.09.242941

**Authors:** A. L. Ayobami, E. A. Kade, K. A. Oladimeji, S. Kehinde, K. Gurpreet

**Affiliations:** Department of Microbiology, University of Lagos (UNILAG), Akoka, Lagos, Nigeria; National University of Ireland Galway, Ireland and University of Limerick; Department of Cell biology and Genetics, University of Lagos (UNILAG), Akoka, Lagos, Nigeria; Department of Health and Health Care Administration, Swami Rama Himalayan University (SRHU), Dehradun, Indian

**Keywords:** Antidiabetic, *Moringa oleifera*, *Ocimum gratissimum Vernonia amygdalina* alloxan-induced

## Abstract

The incidence of diabetes mellitus (DM) is increasing globally and it is a major source of concern. This study was undertaken to assess the antidiabetic effect of the aqueous extract of M*oringa oleifera, Ocimum gratissimum and Vernonia amygdalina*. Sixty adult Wistar rats with body weight of 120-150 g were randomly assigned to groups of five rats each (n=12). Groups 1 served as normal control; Groups 2-5 were diabetic groups; group 2 served as negative control; group 3-5 received 100, 200 and 400 mg/kg of triherbal formula respectively. The body weight (BW) and fasting blood glucose level (FBSL) of the rats were monitored weekly. At the end of the experiment, all the rats were anaesthetized intraperitoneally (I.P) and blood samples were collected by cardiac puncture for biochemical analysis. There was an increase in the BW of the control group and varying doses of tri-herbal formation. It caused 88.0% decrease in FBSL; 371.7%, 386.6% and 296.0% with respect to 100, 200 and 400 mg/kg. Sub-chronic study of the effect of the extract showed a significant increase (P<0.05) in packed cell volume (PCV), white blood counts in rat induced diabetes. The histological studies showed that the diabetic rats with the architecture of the pancreas distorted, was restored to normal by the extract. Its LD50 was found to be greater than 1000 mg/kg indicating its safety in rats. This study has shown that triherbal formula has hypoglycemic and haematogical effects.

## 1. INTRODUCTION

Diabetes mellitus (MD) is characterised by absolute or relative deficiencies in insulin secretion and insulin **a**ction associated with chronic hyperglycemia and disturbances of carbohydrate, lipid and protein metabolism [1-3]. It results from either defect in insulin secretion by the pancreas and inability of the cells to utilize the insulin produced [4-5]. Insulin is secreted by the beta-cells of the islet of Langerhans in the pancreas. Diabetes is a prime risk factor for cardiovascular disease and also affects the heart muscle causing both systolic and diastolic heart failure and also includes polyuria, polyphagia, polydipsia, and ketosis [5].

DM is a global burden, associated with life-threatening complications including stroke, renal failure and cardiac attack. Life for a person with diabetes mellitus means constant awareness of the illness, one or two insulin shots a day, frequent finger punctures to monitor blood glucose level, a restrictive diet, and concern over complications. Global diabetes prevalence has more than doubled over the last three decades, with prevalence rates far exceeding modeled projections; even after allowing for improved surveillance. Nearly 1 in 10 adults worldwide are now affected with diabetes [6-8]

The World Health Organization estimated that over 86% of the people in developing countries rely on traditional remedies such as herbs for their daily needs and about 855 traditional medicines include crude plant extracts (6). DM is one of the world’s leading causes of death, with over 150 million diabetic cases worldwide (9). More than 1.70 million Nigerians above 15 years old are diabetic, with about 70,000 children less than 15 years developing Type 1 diabetes annually. Diabetes is prevalently rising globally as a result of obesity, population growth and sedentary lifestyles, and it is projected to be over 360 million cases by 2030 (25).

Over 50% of plants serve in traditional medicine for their health benefits in combating certain ailments affecting humans such as dysentery, diarrhea, toothache, skin infections and diabetes. Many conventional drugs have been derived from prototypic molecules in medicinal plants. To date, over 400 traditional plant treatments for diabetes have been reported, although only few of the plants have received scientific and medical evaluation to assess their efficacy (11). The attributed hypoglycemic effects of these plants is due to their ability to restore the function of pancreatic tissues by causing an increase in insulin output or inhibit the intestinal absorption of glucose or to the facilitation of metabolites in insulin dependent processes. Hence, treatment with herbal drugs has an effect on protecting β-cells and smoothing out fluctuation in glucose levels. Most of these plants have been found to have chemical constituents like glycosides, alkaloids, phenols, terpenoid and flavonoids that are frequently connected as having antidiabetic effects, (12).

*Moringa oleifera* is the most widely cultivated species of a monogeneric family, the Moringaceae. Native to the sub-Himalayan tracts it is now widely cultivated and has become naturalized in many locations in the tropics [13]. The relative ease with which it propagates through both sexual and asexual means and its low demand for soil nutrients and water after being planted makes its production and management easy [14]. Moringa preparations have been cited in the scientific literature as having nutritional, antimicrobial, hypotensive, antispasmodic, antiulcer, anti-inflammatory, hypo-cholesterolemic, and hypoglycemic activities, as well as having considerable efficacy in water purification by flocculation, sedimentation, antibiosis and even reduction of Schistosome cercariae titer [15]. The plant has also been used extensively for treating inflammation, cardiovascular disease, liver disease and hematological, hepatic and renal function [16]. Moringa leaf extracts have also been shown to have ameliorating effect on chromium-induced testicular toxicity in rats [17] and to enhance sexual activity in mice [18].

*Ocimum gratissimum*, African sweet basil, is a plant belonging to the Lamiceae family. The Ocimum species are widely found in tropical and subtropical regions and commonly used as food spice and traditional herb. The herb has been recommended for the treatment of various diseases [19]. Its hypoglycemic efficacy has been reported [20-21] as has its therapeutic potentials on hepatic disorders [22].

*Vernonia amygdalina*, a member of Asteracea, found throughout the tropical Africa. It is commonly known as bitter leaf. The parts of the plants mostly used are the leaves, usually used in the treatment of diarrhoea, headache, wart and kidney function [23]

## 2. MATERIALS AND METHODS

### 2.1 Collection of Plant Materials

Fresh matured Moringa oleifera, Ocimum gratissimum and Vernonia amygdalina leaves were harvested from the Botanical Garden, University of Lagos, Nigeria and were authenticated at the Department of Botany, University of Lagos, Nigeria. Voucher specimens’ were deposited in the Department of Botany herbarium, University of Lagos. The leaves were rinsed severally with clean tap water to remove dust particles and debris and thereafter allowed to dry completely. About 2kg portion of each leave was then taken for preparation of the plant extracts.

### 2.2 Preparation of Plant Extract

Two kilograms (2kg) each of M. oleifera, O. gratissimum and V. amygdalina was separately homogenised with an electric blender in 4.0 liters of 80% (v/v) ethanol. The blends were allowed for 48hours in a refrigerator at 4ºC to allow for thorough extraction of the plants’ active components. These were then filtered with a cheese cloth and later with Whatman No. 1filter paper to obtain a homogenous filtrate. The filtrates were then concentrated in vacuo at low temperatures (37-40ºC) to about one tenth the original volume using a rotary evaporator. The concentrates were allowed open in a water bath at 40^°^C to allow for complete dryness yielding 80.00g (4.0%), 75.50g (3.8%) and 75. 50g of greenish brown substance obtained.

### 2.3 Acute Toxicity Test

The acute toxicity and lethality test of the aqueous plant extracts in rats was estimated using the method of Lorke as described by (Okoli et al., 24), and the animals were observed continuously after every 2 hrs. Under the following profiles:

**Behavioral Profile**: Alertness, Restlessness

**Neurological Profile:** Pain Response, Touch Response, Gait

**Autonomic Profile:** Defecation and urination

After a period of 24 hrs., the animals were observed for any lethality or death.

### 2.4 Experimental Animals

Sixty adult male albino Wistar rats with a body weight of 120-180g obtained from the animal house of the Department of Biological Sciences of Federal University of Agriculture, Abeokuta, Nigeria were used for this study. The animals were placed in standard cages maintained in 12-hour light: dark cycle under standard conditions (temperature 25+5°C, relative humidity 50+5%). All animals received standard laboratory animal’s chow and water ad libitum during the whole period of experiment.

### 2.5 Experimental Induction of Diabetes

Diabetes was induced according to the principle of Aruna et al., as described by (Okoli et al., 24) with slight modification. A single intraperitoneal injection of alloxan monohydrate (150 mg/kg b.w.) was given and after day 5, blood was drawn from tail vein of the rat by snipping to determine blood glucose level with an automated glucose analyzed device (One Touch Glucometer, Lifecan, California, USA. Animals with blood glucose level ≥ 225 mg/dl were considered diabetic and used for the study.

### 2.6 Experimental Design

Sixty rats were divided into five different experimental groups of 12 animals each. Two groups, normal control group (NC) and diabetic control group (DC) received placebo, 0.5ml DMSO; the other three diabetic groups, the treated groups, received respectively Insulin (administered at a dose of 5Ukg-1b.w.s.c once per day to simulate human regimens), 100mg/kg, 200mg/kg and 400mg/kg of a combination of M. oleifera, O. gratissimum and V. amygdalina. The plant extracts were administered orally. The dosage of the extract was determined from preliminary studies in our laboratory. The duration of treatment was 28 days after induction of diabetes. At the end of the experimental period, the animals were fasted for 12h, then anaesthetized under chloroform vapour and dissected. Whole blood was obtained by cardiac puncture into sterile plain tubes for the hematological assays. The organs were also surgically removed for histological study.

### 2.7 Statistical Analysis

Results were analyzed using SPSS. All data were reported as mean ± SEM. For the evaluation of significant differences, the comparison of means between any two experimental groups was done by computer program for student-t test (paired t-test). Differences were considered to be statistically significant if P = 0.5.

## 3. RESULTS

### 3.1 Body Weight

The extract showed a significant increase (P<0.05) in body weight when compared with pre-diabetic values (Table 1.)

**Table 1:** Effect of M*oringa oleifera, Ocimum gratissimum and Vernonia amygdalina* combination on the body weight of diabetic rats.

| Treatment | BLOOD GLUCOSE LEVEL(mg/dl) |  |  |  |
| --- | --- | --- | --- | --- |
|  | Pre-treatment | Day 7 | Day 14 | Day 28 |
| Control | 87.8 | 103.4* | 122.6* | 156.6* |
| Negative | 90.4 | 106.4* | 107.2* | 132.8* |
| 100 mg/kg | 74.6 | 81.4* | 98.1* | 109.6* |
| 200 mg/kg | 77.6 | 82.6* | 99.4* | 107.8* |
| 400 mg/kg | 79.5 | 83.9* | 89.8* | 104.6* |

Value express as mean. Values with superscripts\* represent increase in body weight relative to pre-treatment

**Figure 1:**
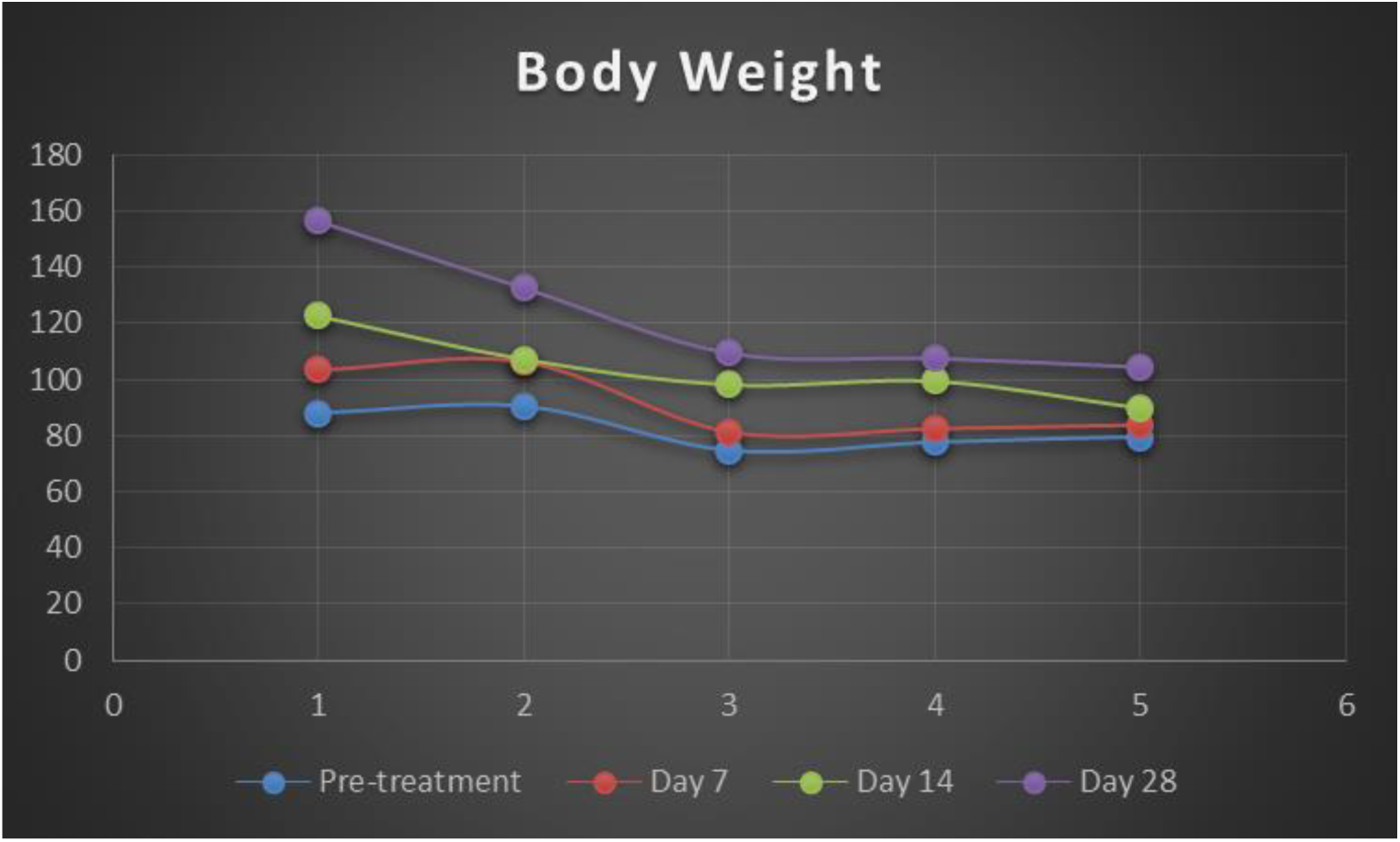
Effect of M*oringa oleifera, Ocimum gratissimum and Vernonia amygdalina* combination on the body weight of diabetic rats.

### 3.2. Anti-Diabetic Effect of of M*oringa oleifera, Ocimum gratissimum and Vernonia amygdalina* combination

The extract showed a significant reduction (P<0.05) in blood glucose levels following sub-chronic administration when compared with the diabetic pre-treatment values. At day 28, the extract reduced the blood glucose level (Table 2).

**Table 2:**
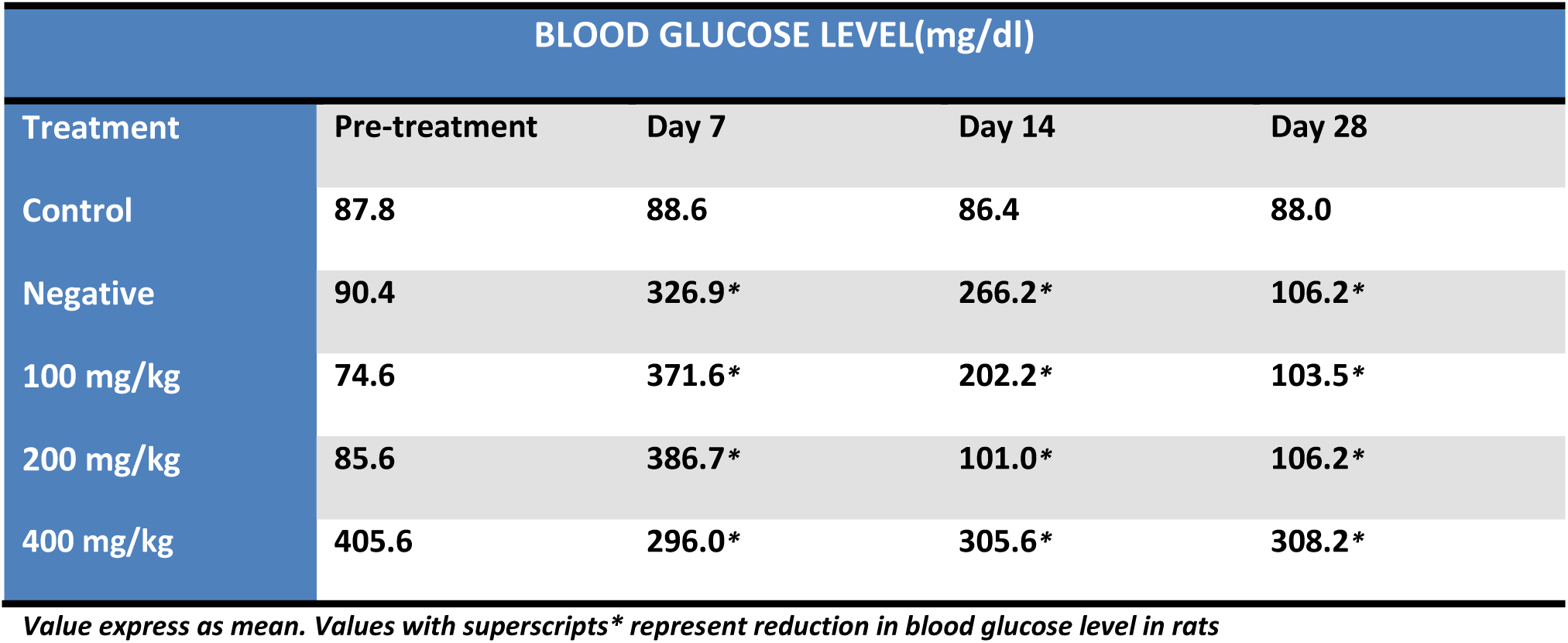
Effect of M*oringa oleifera, Ocimum gratissimum and Vernonia amygdalina* combination on blood glucose level of diabetic rats.

### 3.3 Haematological Parameters

The extract showed a significant reduction (P<0.05) in blood haematological parameters when compared with the diabetic pre-treatment values (Table 3).

**Table 3:**
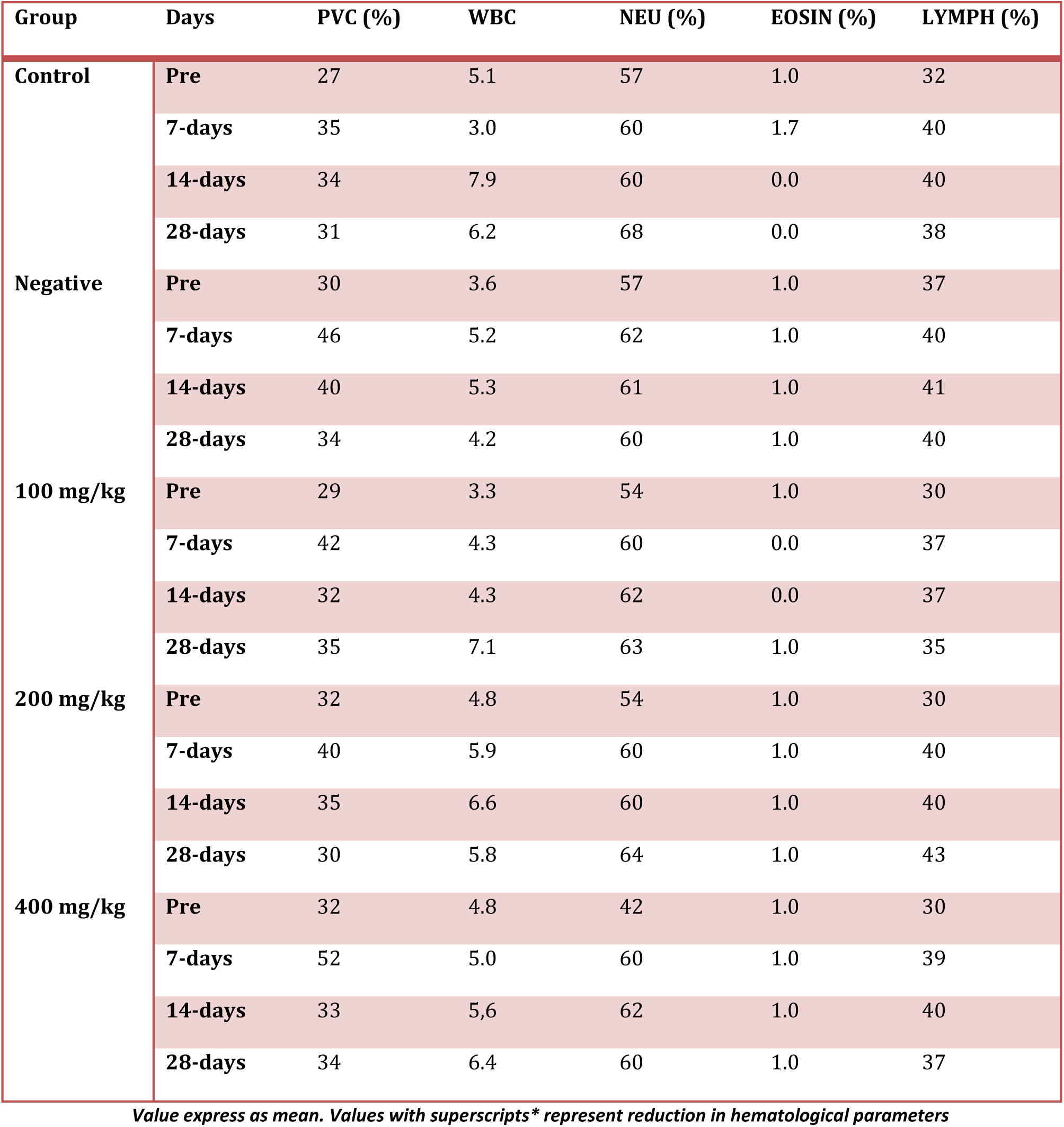
Effect of M*oringa oleifera, Ocimum gratissimum and Vernonia amygdalina* on haematological parameter on diabetic rats.

| Group | Days | PVC (%) | WBC | NEU (%) | EOSIN (%) | LYMPH (%) |
| --- | --- | --- | --- | --- | --- | --- |
| Control | Pre | 27 | 5.1 | 57 | 1.0 | 32 |
|  | 7-days | 35 | 3.0 | 60 | 1.7 | 40 |
|  | 14-days | 34 | 7.9 | 60 | 0.0 | 40 |
|  | 28-days | 31 | 6.2 | 68 | 0.0 | 38 |
| Negative | Pre | 30 | 3.6 | 57 | 1.0 | 37 |
|  | 7-days | 46 | 5.2 | 62 | 1.0 | 40 |
|  | 14-days | 40 | 5.3 | 61 | 1.0 | 41 |
|  | 28-days | 34 | 4.2 | 60 | 1.0 | 40 |
| 100 mg/kg | Pre | 29 | 3.3 | 54 | 1.0 | 30 |
|  | 7-days | 42 | 4.3 | 60 | 0.0 | 37 |
|  | 14-days | 32 | 4.3 | 62 | 0.0 | 37 |
|  | 28-days | 35 | 7.1 | 63 | 1.0 | 35 |
| 200 mg/kg | Pre | 32 | 4.8 | 54 | 1.0 | 30 |
|  | 7-days | 40 | 5.9 | 60 | 1.0 | 40 |
|  | 14-days | 35 | 6.6 | 60 | 1.0 | 40 |
|  | 28-days | 30 | 5.8 | 64 | 1.0 | 43 |
| 400 mg/kg | Pre | 32 | 4.8 | 42 | 1.0 | 30 |
|  | 7-days | 52 | 5.0 | 60 | 1.0 | 39 |
|  | 14-days | 33 | 5.6 | 62 | 1.0 | 40 |
|  | 28-days | 34 | 6.4 | 60 | 1.0 | 37 |

*Value express as mean. Values with superscripts\* represent reduction in hematological parameters*

### 3.4 Histological Studies

The effects of the extract on pancreatic tissues of diabetic-treated were shown in plates 1-5.

**Plate 1:**
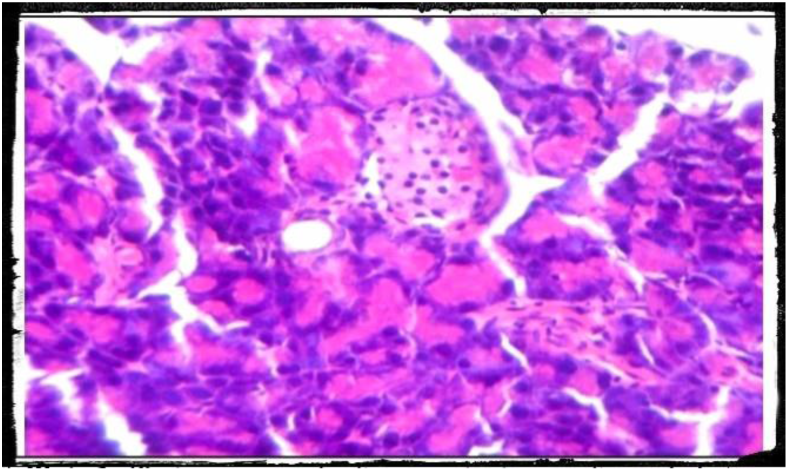
Photomicrography of pancreatic tissue of control (H and E x400)

**Plate 2:**
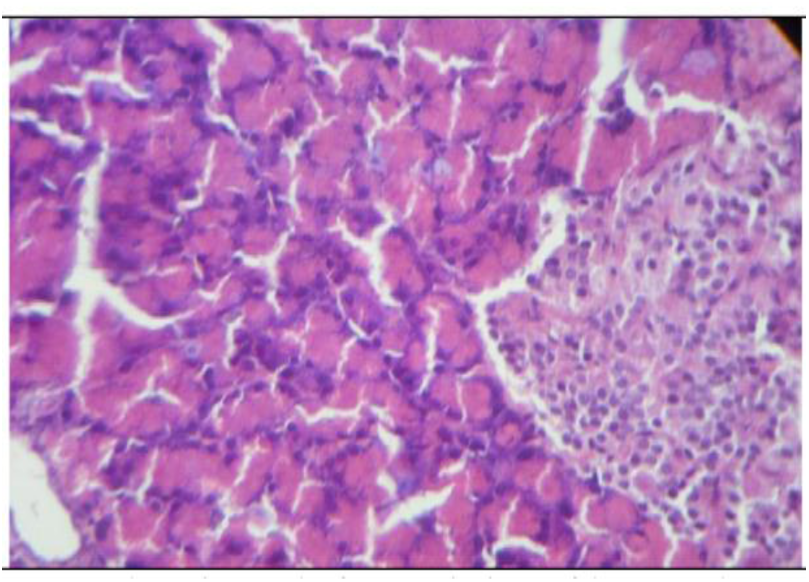
Photomicrography of pancreatic tissue of control (H and E x400). There was no visible change in the structure.

**Plate 3:**
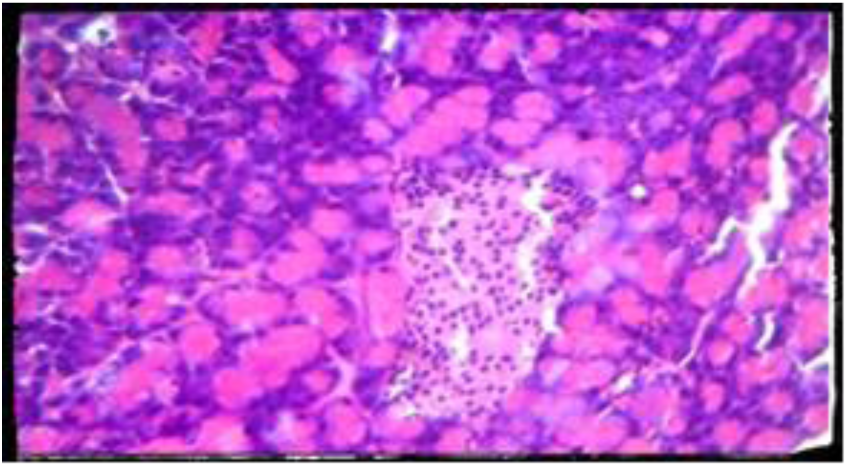
Photomicrography of pancreatic tissue of control (H and E x400). There was no visible change in the pancreas of animals administered with 100 mg/kg compare to control.

**Plate 4:**
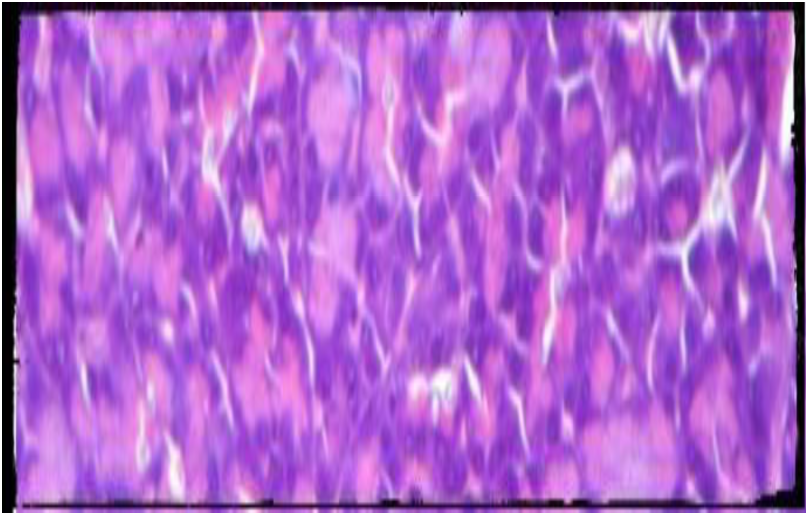
Photomicrography of pancreatic tissue of control (H and E x400). There was no visible change in the pancreas of animals administered with 200 mg/kg compare to control.

**Plate 5:**
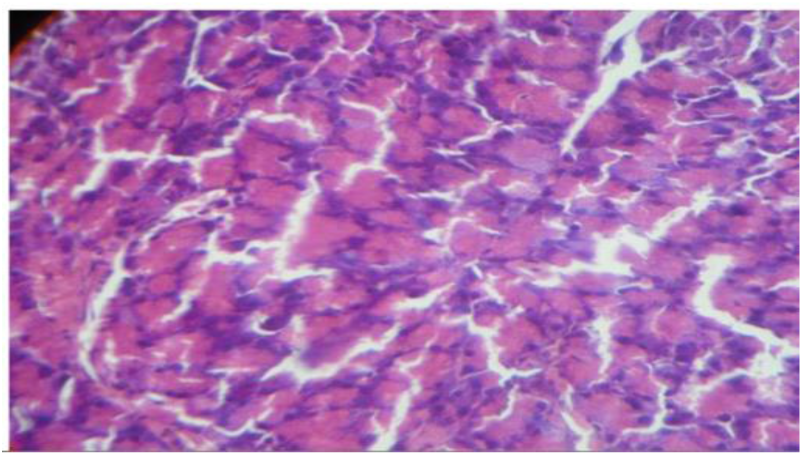
Photomicrography of pancreatic tissue of control (H and E x400). There was no visible change in the pancreas of animals administered with 400 mg/kg compare to control.

## 5. Discussion

Diabetes Mellitus (DM) poses a substantial threat to the health and quality of life of an individual. Due to increase complications and mortality in DM, adequate glyceamic controls might be useful in its management. In this study, anti-diabetic potential of aqueous extract of M*oringa oleifera, Ocimum gratissimum and Vernonia amygdalina* in alloxan-induced diabetic rats showed that a single oral administration of the extract reduced the fasting blood glucose levels as well as suppressed the rise in blood glucose of normal rats after a heavy glucose meal. A number of investigators have shown that coumarin, flavonoid, terpenoid and a host of other secondary plant metabolites including arginine and glutamic acids posses’ hypoglycemic effects in various experimental animals model (11). However, in line with the hypothesis of Marles and Farnsworth (18) which stipulates that plant which contains terpenoid posseses hypoglycemic activities in diabetic and normal mammal, then it could be thus concluded that aqueous of M*oringa oleifera, Ocimum gratissimum and Vernonia amygdalina* extract possess hypoglycemic activity.

In the anti-diabetic activity studies, daily oral administration of the extract for 28 days revealed a gradual but sustained reduction in blood glucose levels in diabetic rats. According to Oyedepo et al. 22, aqueous extract of of M*oringa oleifera, Ocimum gratissimum and Vernonia amygdalina* combination has been shown to normalize the high blood glucose level in diabetic rats by the 28th day of the experiment. Treatment with the extract also reduced mortality of diabetic rats from hyperglycaemia and prolonged their survival. In this study, the control animals all died on day 14 post-induction of diabetes (data not shown) whereas the extract-treated group survived beyond the period of the experiment. Since a better activity has been achieved in severely diabetic rats with damaged islet, it is possible that aqueous extract of of M*oringa oleifera, Ocimum gratissimum and Vernonia amygdalina* combination has some direct effect. The increase in the total cholesterol and TAG and the decrease in HDL levels of diabetic rats observed in this study are in accordance with earlier report in diabetic subjects (5). Diabetes-induced hyperlipidemia has been attributed to excess mobilization of fat from the adipose due to underutilization of glucose (9). Chronic oral administration of the extract also revealed a significant reduction in cholesterol levels as well as the increase in HDL levels of the diabetic-treated rats when compared with the control rats supports the findings of Coon and Ernst (8) who stated that most hypoglycemic plants have potentials blood glucose level.

Also significant advancement in the haematological parameters on a long term treatment with extract for 28 days suggests the favourable effects of the aqueous extract of of M*oringa oleifera, Ocimum gratissimum and Vernonia amygdalina* combination. Sub-chronic administration of the extract also showed weight gain in diabetic-treated compared to control rats. Histopathological analysis showed that after the long administration of the extract at 100, 200 and 400 mg/kg body weight, the extract was able to restore the structure of the pancreas damaged by alloxan. (Plates 1-5). Alloxan owes its diabetogenic potential to destruction of β -cells of the islet (12-13) which consequently impairs insulin secretion and leads to hyperglycaemia. Extract treatment may have restored the integrity and possibly, functions of the damaged pancreatic tissues. The precise mechanism of this tissue repair is yet to be known. However, due to the tremendous implication of oxidative stress (12-13) which leads to the damage to the pancreas, it seems reasonable to suggest that the antioxidant (27) and radical scavenging effects (14) of Moringa oleifera leaf may play a significant role in protecting pancreatic tissues from oxidants including those generated by alloxan. Alloxan destroys insulin-producing pancreatic β-cells through the production of reactive oxygen species which then cause tissue damage (16). The hypoglycaemic effect of the extract may be responsible for protection confer on the pancreas from the deleterious effect of chronic hyperglycaemia. Rather than reflecting a direct tissue repair effect, it is likely that the extract, through antioxidative and hypoglycaemic effects, protected the already compromised pancreas from further assault or pancreatic damage which may allow the natural repair processes to proceed and restore the tissues. However, it is unknown if the repaired tissues also had their functions fully or partially restored since the blood glucose level of the animals did not return to normalcy or pre-treatment levels as at the end of the experiment. A return to pre-treatment levels might indicate a full restoration of insulin secretion by the repaired pancreatic tissues (20).

## Ethical Approval

All authors hereby declare that the research has been determine exempt from review by the university animal research and ethics review committee and that the principles of the laboratory animal care were followed

## Acknowledgements

This work was supported by students and provided by Department of cell biology and Genetics. The authors are also grateful to Lagos State University Teaching Hospital for the histological analyses.

